# Three new genome assemblies support a rapid radiation in *Musa acuminata* (wild banana)

**DOI:** 10.1101/306605

**Authors:** M Rouard, G Droc, G Martin, J Sardos, Y Hueber, V Guignon, A Cenci, B Geigle, M S Hibbins, N Yahiaoui, F-C Baurens, V Berry, M W Hahn, A D’Hont, N Roux

**Author notes:** Corresponding author: Mathieu Rouard.

## Abstract

Edible bananas result from interspecific hybridization between *Musa acuminata* and *Musa balbisiana*, as well as among subspecies in *M. acuminata*. Four particular *M. acuminata* subspecies have been proposed as the main contributors of edible bananas, all of which radiated in a short period of time in southeastern Asia. Clarifying the evolution of these lineages at a whole-genome scale is therefore an important step toward understanding the domestication and diversification of this crop. This study reports the *de novo* genome assembly and gene annotation of a representative genotype from three different subspecies of *M. acuminata*. These data are combined with the previously published genome of the fourth subspecies to investigate phylogenetic relationships and genome evolution. Analyses of shared and unique gene families reveal that the four subspecies are quite homogenous, with a core genome representing at least 50% of all genes and very few *M. acuminata* species-specific gene families. Multiple alignments indicate high sequence identity between homologous single copy-genes, supporting the close relationships of these lineages. Interestingly, phylogenomic analyses demonstrate high levels of gene tree discordance, due to both incomplete lineage sorting and introgression. This pattern suggests rapid radiation within *Musa acuminata* subspecies that occurred after the divergence with *M. balbisiana*. Introgression between *M. a.* ssp. *malaccensis* and *M. a.* ssp. *burmannica* was detected across a substantial portion of the genome, though multiple approaches to resolve the subspecies tree converged on the same topology. To support future evolutionary and functional analyses, we introduce the PanMusa database, which enables researchers to exploration of individual gene families and trees.

## Background

Bananas are among the most important staple crops cultivated worldwide in both the tropics and subtropics. The wild ancestors of bananas are native to the Malesian Region (including Malaysia and Indonesia) (Simmonds 1962) or to northern Indo-Burma (southwest China). Dating back to the early Eocene (Janssens et al. 2016), the genus *Musa* currently comprises 60 to 70 species divided into two sections, Musa and Callimusa (Häkkinen 2013). Most of modern cultivated bananas originated from natural hybridization between two species from the section Musa, *Musa acuminata*, which occurs throughout the whole southeast Asia region, and *Musa balbisiana*, which is constrained to an area going from east India to south China (Simmonds & Shepherd 1955). While no subspecies have been defined so far in *M. balbisiana*, *M. acuminata* is further divided into multiple subspecies, among which at least four have been identified as contributors to the cultivated banana varieties, namely *banksii, zebrina, burmannica*, and *malaccensis* (reviewed in Perrier et al. 2011). These subspecies can be found in geographical areas that are mostly non-overlapping. *Musa acuminata* ssp. *banksii* is endemic to New Guinea. *M. a*. ssp. *zebrina* is found in Indonesia (Java island), *M. a.* ssp. *malaccensis* originally came from the Malay Peninsula (De Langhe et al. 2009; Perrier et al. 2011), while *M. a*. ssp. *burmannica* is from Burma (today’s Myanmar) (Cheesman 1948).

While there are many morphological characters that differentiate *M. acuminata* from *M. balbisiana*, the subspecies of *M. acuminata* have only a few morphological differences between them. For instance, *M. a*. ssp. *burmannica* is distinguished by its yellowish and waxless foliage, light brown markings on the pseudostem, and by its compact pendulous bunch and strongly imbricated purple bracts. *M. a*. ssp. *banksii* exhibits slightly waxy leaf, predominantly brown-blackish pseudostems, large bunches with splayed fruits, and non-imbricated yellow bracts. *M. a*. ssp. *maaccensis* is strongly waxy with a horizontal bunch, and bright red non-imbricated bracts, while *M. a*. ssp. *zebrina* is characterized by dark red patches on its dark green leaves (Simmonds 1956).

Previous studies based on a limited number of markers have been able to shed some light on the relationships among *M. acuminata* subspecies (Sardos et al. 2016; Christelová et al. 2017). Phylogenetic studies have been assisted by the availability of the reference genome sequence for a representative of *M. acuminata* ssp. *malaccensis* (D’Hont et al. 2012; Martin et al. 2016) and a draft *M. balbisiana* genome sequence (Davey et al. 2013). However, the availability of large genomic datasets from multiple (sub)species are expected to improve the resolution of phylogenetic analyses, and thus to provide additional insights on species evolution and their specific traits (Bravo et al. 2018). This is especially true in groups where different segments of the genome have different evolutionary histories, as has been found in *Musaceae* (Christelová, Valárik, et al. 2011). Whole-genome analyses also make it much easier to distinguish among the possible causes of gene tree heterogeneity, especially incomplete lineage sorting (ILS) and hybridization (Folk et al. 2018).

Moreover, the availability of multiple reference genome sequences opens the way to so-called pangenome analyses, a concept coined by Tettelin et al. (2005). The pangenome is defined as the set of all gene families found among a set of phylogenetic lineages. It includes i) the core genome, which is the pool of genes common to all lineages, ii) the accessory genome, composed of genes absent in some lineages, and iii) the species-specific or individual-specific genome, formed by genes that are present in only a single lineage. Identifying specific compartments of the pangenome (such as the accessory genome) offers a way to detect important genetic differences that underlie molecular diversity and phenotypic variation (Morgante et al. 2007).

Here, we generated three *de novo* genomes for the subspecies *banksii, zebrina and burmannica*, and combined these with existing genomes for *M. acuminata* ssp. *malaccensis* (D’Hont et al. 2012) and *M. balbisiana* (Davey et al. 2013). We thus analyzed the whole genome sequences of five extant genotypes comprising the four cultivated bananas’ contributors from *M. acuminata*, i.e. the reference genome ‘DH Pahang’ belonging to *M. acuminata* ssp. *malaccensis*, ‘Banksii’ from *M. acuminata* ssp. *banksii*, ‘Maia Oa’ belonging to *M. acuminata* ssp. *zebrina*, and ‘Calcutta 4’ from *M. acuminata* ssp. *burmannica*, as well as *M. balbisiana* (i.e. ‘Pisang Klutuk Wulung’ or PKW). We carried out phylogenomic analyses that provided evolutionary insights into both the relationships and genomic changes among lineages in this clade. Finally, we developed a banana species-specific database to support the larger community interested in crop improvement.

## Results

### Assembly and gene annotation

We generated three *de novo* assemblies belonging to *M. acuminata* ssp. *banksii*, *M. a*. ssp. *zebrina* and *M. a*. ssp. *burmannica* (**S Table 1 & 2**). The number of predicted protein coding genes per genome within different genomes of *Musa* ranges from 32,692 to 45,069 (**S Table 4**). Gene number was similar for *M. a*. ssp. *maaccensis* ‘DH Pahang’, *M. balbisiana* ‘PKW’ and *M. a*. ssp. *banksii* ‘Banksii’ but higher in *M. a*. ssp. *zebrina* ‘Maia Oa’ and *M. a*. ssp. *burmannica* ‘Calcutta 4’. According to BUSCO (**S. Table 3**), the most complete gene annotations are ‘DH Pahang’ (96.5%), ‘Calcutta 4’ (74.2%) and ‘Banksii’ (72.5%), followed by ‘PKW’ (66.5%) and ‘Maia Oa’ (61.2%).

### Gene families

The percentage of genes in orthogroups (OGs), which is a set of orthologs and recent paralogs (*i.e*. gene family), ranges from 74 in *M. a*. *zebrina* ‘Maia Oa’ to 89.3 in *M a*. *malaccensis* ‘DH Pahang’ with an average of 79.8 (Table 1). Orthogroups have a median size of 4 genes and do not exceed 50 (**S. Table5**). A pangenome here was defined on the basis of the analysis of OGs in order to define the 1) core, 2) accessory, and 3) unique gene set(s). On the basis of the five genomes studied here, the pangenome embeds a total of 32,372 OGs composed of 155,222 genes. The core genome is composed of 12,916 OGs (Figure 1). Among these, 8,030 are composed of only one sequence in each lineage (*i.e*. are likely single-copy orthologs). A set of 1489 OGs are specific to all subspecies in *M. acuminata*, while the number of genes specific to each subspecies ranged from 14 in the *M. acuminata* ‘DH Pahang’ to 110 in *M. acuminata* ‘Banksii’ for a total of 272 genes across all genotypes. No significant enrichment for any Gene Ontology (GO) category was detected for subspecies-specific OGs (**S. data 1**).

**Figure 1.**
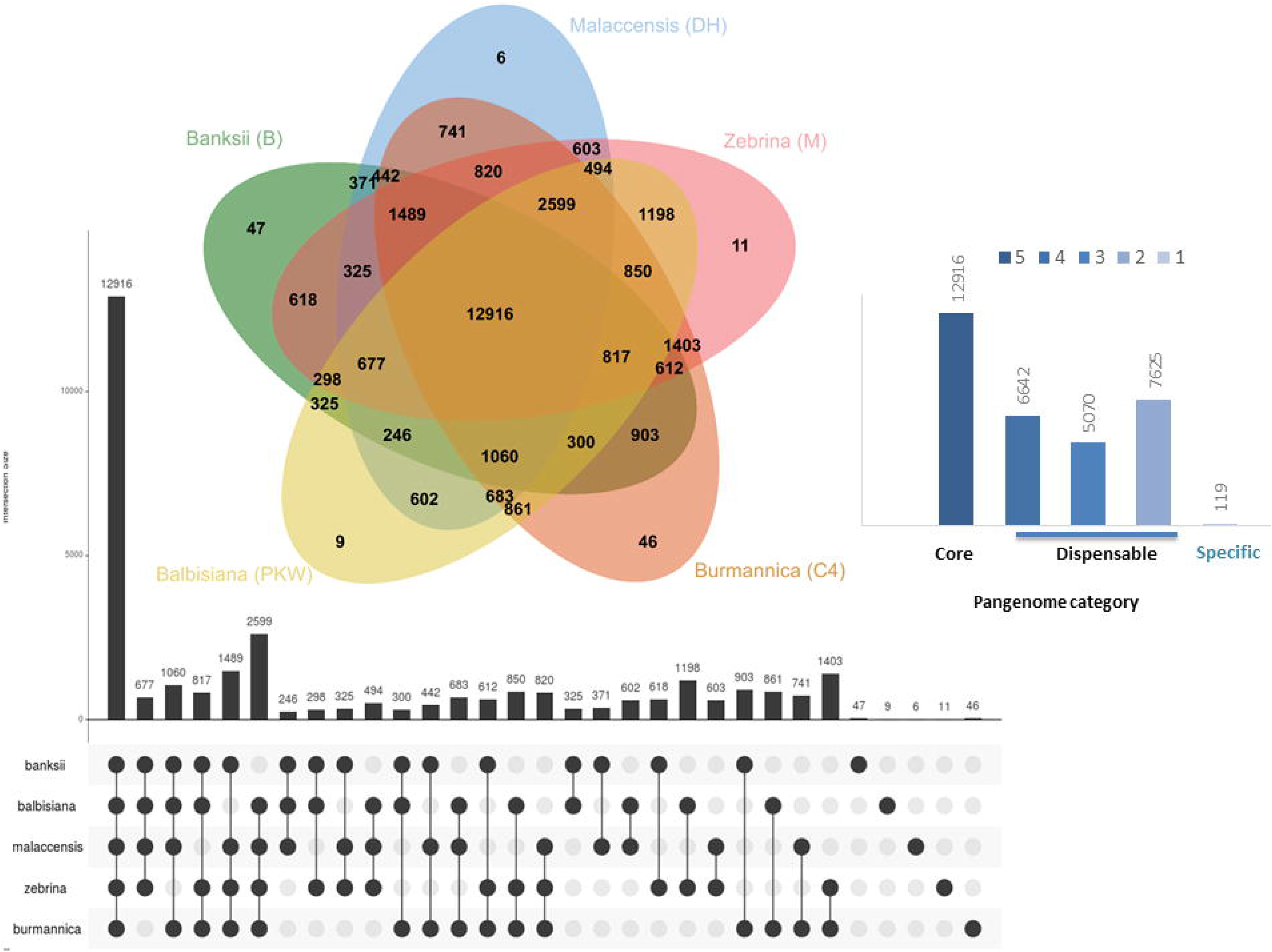
Five-way Venn diagram showing the distribution of shared gene families. (at least two sequences per OG) among *M. a. banksii* ‘Banksii’, *M. a. zebrina* ‘Maia Oa’, *M. a. burmannica* ‘Calcutta 4’, *M. a. malaccensis* ‘DH Pahang’ and *M. balbisiana* ‘PKW’ genomes. On the right, number of orthologous groups by species and pangenome category. At the bottom, same dataset visualized with UpsetR (Lex et al. 2014).

**Table 1.** Summary of the gene clustering statistics per (sub)species.

|  | *M.*<br>*acuminata*<br>*malaccensis*<br>‘DH<br>Pahang’ | *M.*<br>*acuminata*<br>*burmannica*<br>‘Calcutta 4’ | *M.*<br>*acuminata*<br>*banksii*<br>‘Banksii’ | *M.*<br>*acuminata*<br>*zebrina*<br>‘Maia<br>Oa’ | *M.*<br>*balbisiana*<br>‘PKW’ |
| --- | --- | --- | --- | --- | --- |
| # genes | 35,276 | 45,069 | 32,692 | 44,702 | 36,836 |
| # genes in orthogroups | 31,501 | 34,947 | 26,490 | 33,059 | 29,225 |
| # unassigned genes | 3,775 | 10,122 | 6,202 | 11,643 | 7,611 |
| % genes in orthogroups | 89.3 | 77.5 | 81 | 74 | 79.3 |
| % unassigned genes | 10.7 | 22.5 | 19 | 26 | 20.7 |
| # orthogroups containing species | 24,074 | 26,542 | 21,446 | 25,730 | 23,935 |
| % orthogroups containing species | 74.4 | 82 | 66.2 | 79.5 | 73.9 |
| # species-specific orthogroups | 6 | 46 | 47 | 11 | 9 |
| # genes in species-specific orthogroups | 14 | 104 | 110 | 23 | 21 |
| % genes in species-specific orthogroups | 0 | 0.2 | 0.3 | 0.1 | 0.1 |

### Variation in gene tree topologies

Phylogenetic reconstruction performed with single-copy genes (*n*=8,030) showed high levels of discordance among the different individual gene trees obtained, both at the nucleic acid and protein levels (Figure 2A). Considering *M. balbisiana* as outgroup, there are 15 possible bifurcating tree topologies relating the four *M. acuminata* subspecies. For all three partitions of the data - protein, CDS, and gene (including introns and UTRs) - we observed all 15 different topologies (Table 2). We also examined topologies at loci that had bootstrap support greater than 90 for all nodes, also finding all 15 different topologies (Table 2). Among trees constructed from whole genes, topologies ranged in frequency from 13.12% for the most common tree to 1.92% for the least common tree (Table 2) with an average length of the 1342 aligned nucleotide sites for CDS and 483 aligned sites for proteins. Based on these results, gene tree frequencies were used to calculate concordance factors on the most frequent CDS gene trees (Table 2), demonstrating that no split was supported by more than 30% of gene trees (Figure 2B). Therefore, in order to further gain insight into the subspecies phylogeny, we used a combination of different approaches described in the next section.

**Figure 2.**
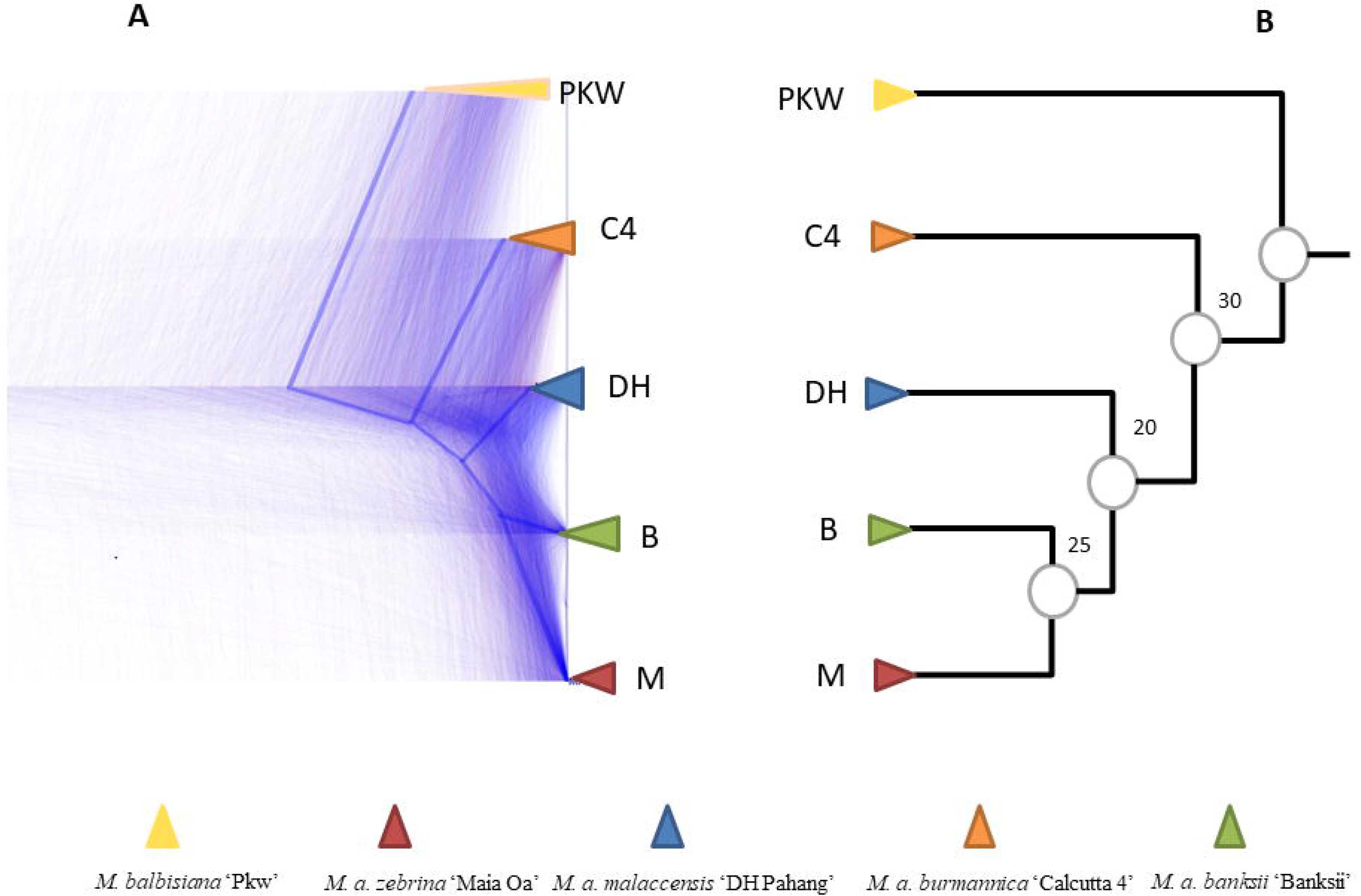
Illustration of gene tree discordance. A) Cloudogram of single copy OGs (CDS) visualized with Densitree. The blue line represents the consensus tree as provided by Densitree B) Species tree with bootstrap-like support based on corresponding gene tree frequency from Table 2 (denoted topology number 2). (PKW = *M. balbisiana* ‘PKW’, C4 = *M. acuminata burmannica* ‘Calcutta 4, M= *M. acuminata zebrina* ‘Maia Oa’, DH= *M. acuminata malaccensis* “DH Pahang’, B = *M. acuminata banksii* ‘Banksii’)

**Table 2.** Frequency of gene tree topologies of the 8,030 single copy OGs. (PKW = *Musa balbisiana* ‘PKW’, C4 = *Musa acuminata burmannica* ‘Calcutta 4, M= *Musa acuminata zebrina* ‘Maia Oa’, DH= *Musa acuminata malaccensis* “DH Pahang’, B = *Musa acuminata banksii* ‘Banksii’). In bold the most frequent topology.

| No. | Topology | # CDS (%) | # Protein (%) | # Gene (%) | # Gene bootstrap >90 (%) |
| --- | --- | --- | --- | --- | --- |
| 1 | (PKW,(C4,(M,(DH,B)))) | **11.9** | 10.58 | **13.12** | 13.72 |
| 2 | (PKW,(C4,(DH,(B,M)))) | 10.8 | 10.48 | 11.92 | 14.88 |
| 3 | (PKW,((DH,C4),(B,M))) | 9.59 | 7.28 | 12.73 | **17.52** |
| 4 | (PKW,(M,(C4,(DH,B)))) | 9.53 | **12.51** | 7.78 | 5.91 |
| 5 | (PKW,(C4,(B,(DH,M)))) | 8.02 | 7.37 | 8.89 | 8.44 |
| 6 | (PKW,((DH,B),(C4,M))) | 7.67 | 6.55 | 9.16 | 12.56 |
| 7 | (PKW,(M,(B,(DH,C4)))) | 6.66 | 8.21 | 5 | 3.06 |
| 8 | (PKW,(B,(M,(DH,C4)))) | 5.58 | 5.23 | 4.61 | 2.53 |
| 9 | (PKW,(DH,(C4,(B,M)))) | 5.41 | 5.21 | 5.18 | 4.96 |
| 10 | (PKW,(B,(C4,(DH,M)))) | 5.26 | 4.45 | 6.2 | 7.07 |
| 11 | (PKW,(B,(DH,(C4,M)))) | 5.02 | 6.82 | 3.36 | 1.9 |
| 12 | (PKW,(M,(DH,(B,C4)))) | 4.23 | 4.68 | 2.84 | 1.16 |
| 13 | (PKW,((DH,M),(B,C4))) | 4.037 | 3.61 | 4.79 | 5.06 |
| 14 | (PKW,(DH,(B,(C4,M)))) | 3.85 | 4.18 | 2.44 | 0.63 |
| 15 | (PKW,(DH,(M,(B,C4)))) | 2.38 | 2.77 | 1.92 | 0.52 |

### Inference of a species tree

We used three complementary methods to infer phylogenetic relationships among the sampled lineages. First, we concatenated nucleotide sequences from all single-copy genes (totaling 11,668,507 bp). We used PHYML to compute a maximum likelihood tree from this alignment, which, as expected, provided a topology with highly supported nodes (Figure 3A). Note that this topology (denoted topology number 1 in Table 2) is not the same as the one previously proposed in the literature (denoted topology number 7 in Table 2) (**S. Figure 1 & 2**).

**Figure 3.**
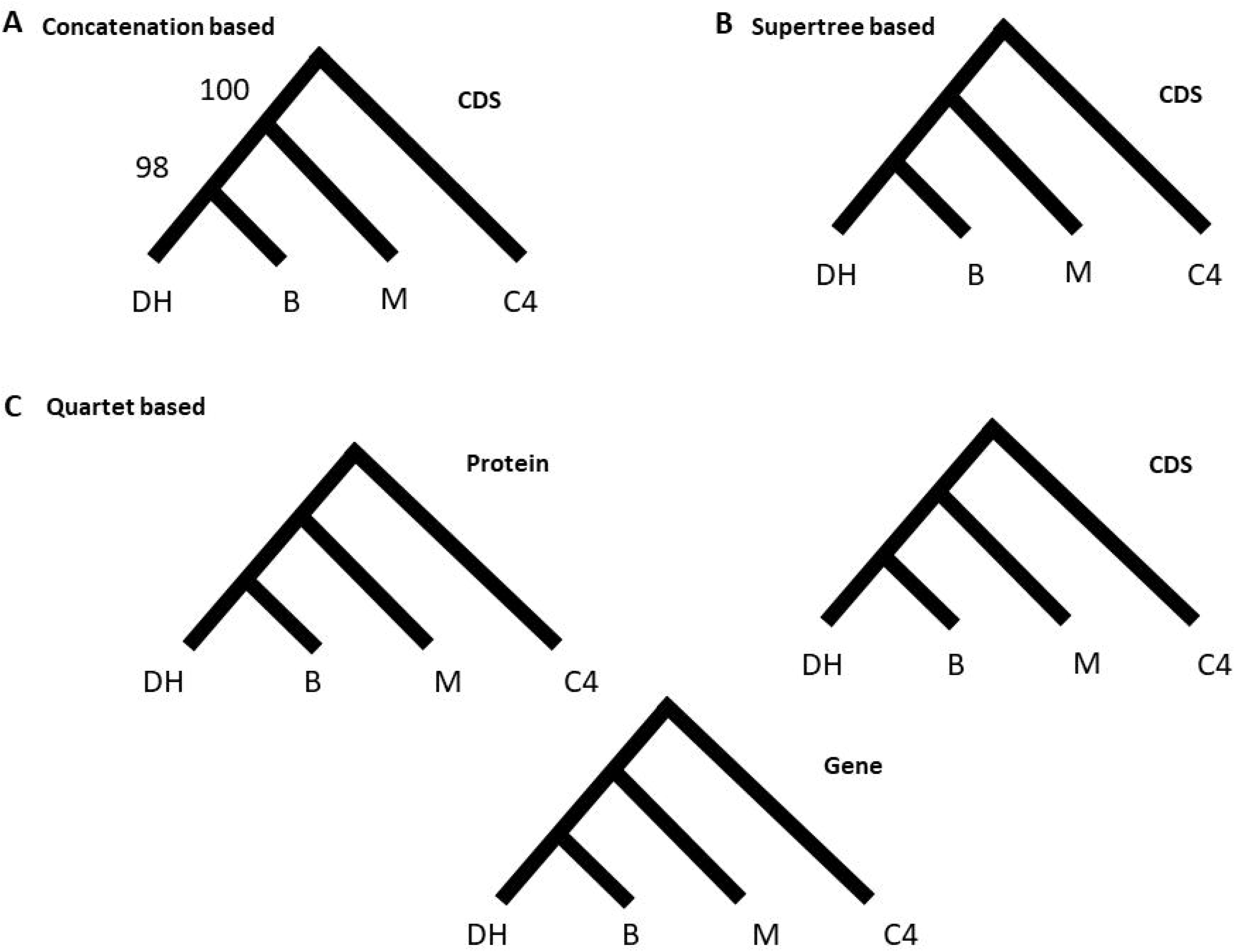
Species topologies computed with three different approaches. A) Maximum likelihood tree inferred from a concatenated alignment of single-copy genes (CDS). B) Supertree-based method applied to single and multi-labelled gene trees C) Quartet-based model applied to protein, CDS, and gene alignments.

Next, we used a method explicitly based on individual gene tree topologies. ASTRAL (Mirarab & Warnow 2015) infers the species tree by using quartet frequencies found in gene trees. It is suitable for large datasets and was highlighted as one of the best methods to address challenging topologies with short internal branches and high levels of discordance (Shi & Yang 2017). ASTRAL found the same topology using ML gene trees from single-copy genes obtained from protein sequences, CDSs, and genes (Figure 3C).

Finally, we ran a supertree approach implemented in PhySIC_IST (Scornavacca et al. 2008) on the single-copy genes and obtained again the same topology (Figure 3B). PhySIC_IST first collapses poorly supported branches of the gene trees into polytomies, as well as conflicting branches of the gene trees that are only present in a small minority of the trees; it then searches for the most resolved supertree that does not contradict the signal present in the gene trees nor contains topological signal absent from those trees. Deeper investigation of the results revealed that ~ 66% of the trees were unresolved, 33% discarded (pruned or incorrectly rooted), and therefore that the inference relied on fewer than 1% of the trees. Aiming to increase the number of genes used by PhySIC_IST, we included multi-copy OGs of the core genome, as well as some OGs in the accessory genomes using the pipeline SSIMUL (Scornavacca et al. 2011). SSIMUL translates multi-labeled gene trees (MUL-trees) into trees having a single copy of each gene (X-trees), i.e. the type of tree usually expected in supertree inference. To do so, all individual gene trees were constructed on CDSs from OGs with at least 4 *M. acuminata* and *M. balbisiana* genes (n=18,069). SSIMUL first removed identical subtrees resulting from a duplication node in these trees, it then filtered out trees where duplicated parts induced contradictory rooted triples, keeping only coherent trees. These trees can then be turned into trees containing a single copy of each gene, either by pruning the smallest subtrees under each duplication node (leaving only orthologous nodes in the tree), or by extracting the topological signal induced by orthology nodes into a rooted triplet set, that is then turned back into an equivalent X-tree. Here we chose to use the pruning method to generate a dataset to be further analyzed with PhySIC_IST, which lead to a subset of 14,507 gene trees representing 44% of the total number of OGs and an increase of 80% compared to the 8,030 single-copy OGs. This analysis returned a consensus gene tree with the same topology as both of the previous methods used here (Figure 3B).

### Evidence for introgression

Although much of the discordance we observe is likely due to incomplete lineage sorting, we also tested for introgression between subspecies. The ABBA-BABA test (Green et al. 2010) was conducted to detect an excess of either ABBA or BABA sites (where “A” corresponds to the ancestral allele and “B” corresponds to the derived allele state) in a four-taxon phylogeny including three *M. acuminata* subspecies as ingroups and *M. balbisiana* as outgroup. Because there were five taxa to be tested, analyses were done with permutation of taxa denoted P1, P2 and P3 and Outgroup (Table 3). Under the null hypothesis of ILS, an equal number of ABBA and BABA sites are expected. However, we always found an excess of sites grouping *malaccensis* (‘DH’) and *burmannica* (‘C4’) (Table 3). This indicates a history of introgression between these two lineages.

**Table 3.** Four-taxon ABBA-BABA Test (*D*-statistic) used for introgression inference from the well-supported topology from Figure 3. ^a^ Discordance = (ABBA + BABA) / Total ^b^ *D* = (ABBA-BABA) / (ABBA+BABA)

| P1 | P2 | P3 | BBAA | ABBA | BABA | Discordance<sup>a</sup> | *D*<sup>b</sup> | P-value |
| --- | --- | --- | --- | --- | --- | --- | --- | --- |
| Malaccensis (DH) | Banksii (B) | Burmannica (C4) | 12185 | 4289 | 8532 | 0.51 | 0.33 | 0 |
| Malaccensis (DH) | Zebrina (M) | Burmannica (C4) | 9622 | 5400 | 9241 | 0.6 | 0.26 | 0 |
| Zebrina (M) | Banksii (B) | Burmannica (C4) | 11204 | 6859 | 6782 | 0.54 | -0.005 | 0.5 |
| Malaccensis (DH) | Banksii (B) | Zebrina (M) | 10450 | 7119 | 6965 | 0.57 | -0.02 | 0.19 |

To test the direction of introgression, we applied the *D*_2_ test (Hibbins and Hahn, unpublished). While introgression between a pair of species (e.g. *malaccensis* and *burmannica*) always results in smaller genetic distances between them, the *D*_2_ test is based on the idea that gene flow in the two alternative directions can also result in a change in genetic distance to other taxa not involved in the exchange (in this case, *banksii*). We computed the genetic distance between *banksii* and *burmannica* in gene trees where *malaccensis* and *banksii* are sister (denoted d_AC_|A,B) and the genetic distance between *banksii* and *burmannica* in gene trees where *malaccensis* and *burmannica* are sister (denoted d_AC_|B,C). The test takes into account the genetic distance between the species not involved in the introgression (*banksii*) and the species involved in introgression that it is not most closely related to (*burmannica*). We identified 1454 and 281 gene trees with d_AC_|A,B=1.15 and d_AC_|B,C = 0.91, respectively, giving a significant positive value of *D*_2_=0.23 (P<0.001 by permutation). These results support introgression from *malaccensis* into *burmannica*, though they do not exclude the presence of a lesser level of gene flow in the other direction.

### PanMusa, a database to explore individual OGs

Since genes underlie traits and wild banana species showed a high level of incongruent gene tree topologies, access to a repertoire of individual gene trees is important. This was the rationale for constructing a database that provides access to gene families and individual gene family trees in *M. acuminata* and *M. balbisiana*. A set of web interfaces are available to navigate OGs that have been functionally annotated using GreenPhyl comparative genomics database (Rouard et al. 2011). Pan*Musa* shares most of the features available on GreenPhyl to display or export sequences, InterPro assignments, sequence alignments, and gene trees (Figure 4). In addition, new visualization tools were implemented, such as MSAViewer (Yachdav et al. 2016) and PhyD3 (Kreft et al. 2017) to view gene trees.

**Figure 4.**
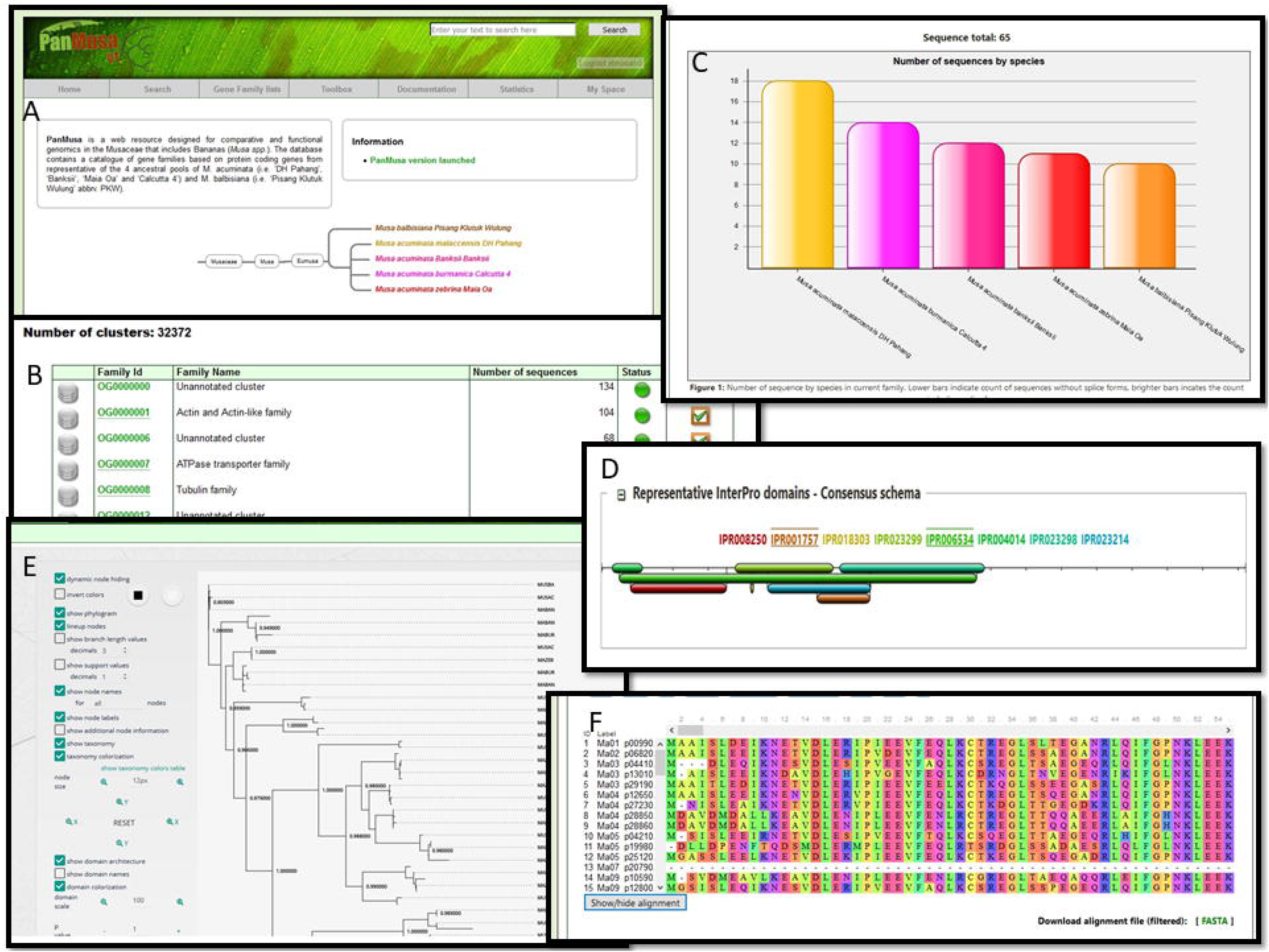
Overview of available interfaces for the PanMusa database. A. Homepage of the website. B. List of functionally annotated OGs. C. Graphical representation of the number of sequence by species. D. Consensus InterPro domain schema by OG. E. Individual gene trees visualized with PhyD3. F. Multiple alignment of OG with MSAviewer.

**Figure 5.**
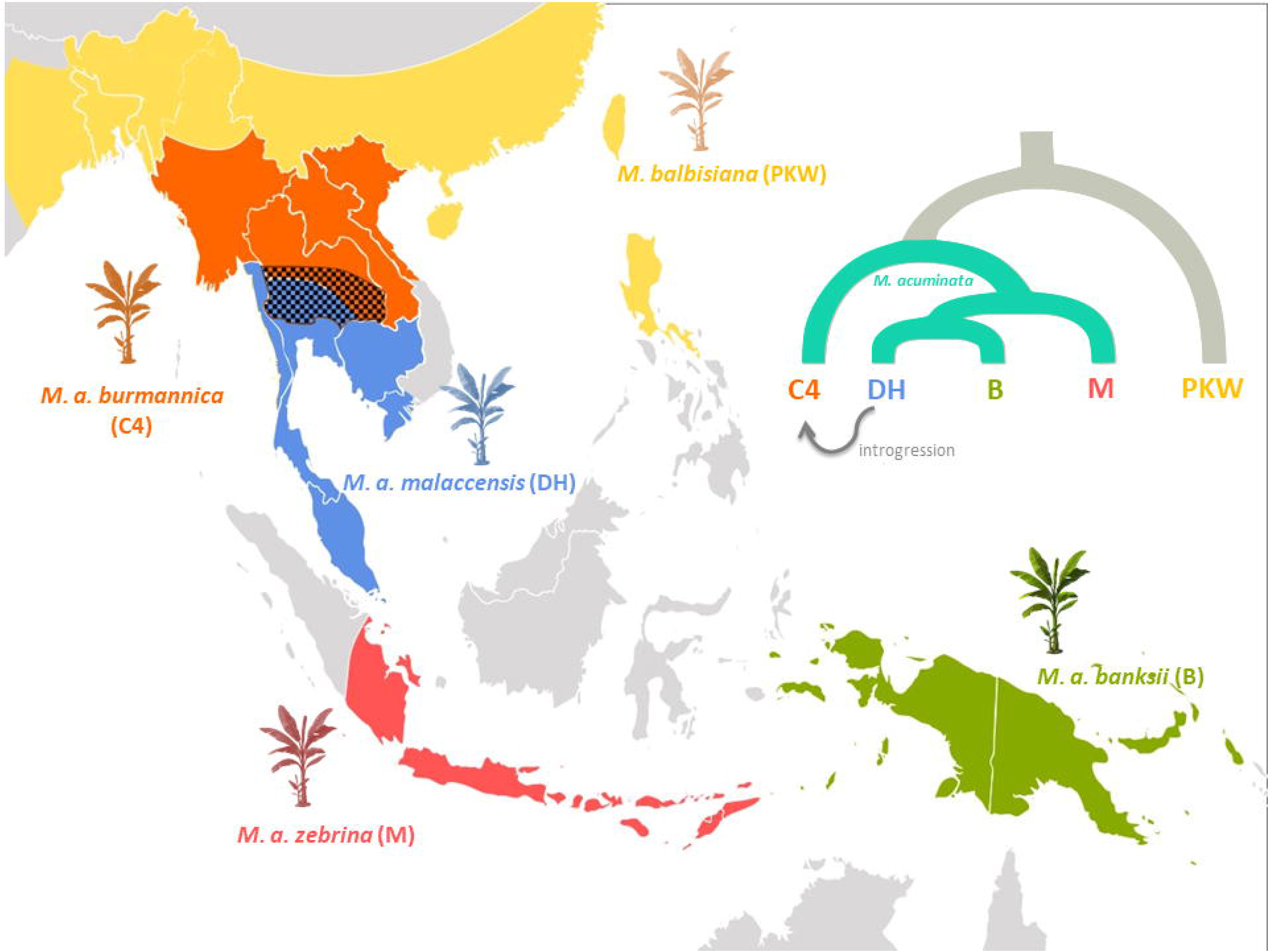
Area of distribution of Musa species in Southeast Asia as described by Perrier et al, 2011; including species tree of *Musa acuminata* subspecies based on results described in Figure 4. Areas of distribution are approximately represented by colors; hatched zone shows area of overlap between two subspecies where introgression may have occurred.

## Discussion

### *M. acuminata* subspecies contain few subspecies-specific families

In this study, we used a *de novo* approach to generate additional reference genomes for the three subspecies of *Musa acuminata*; all three are thought to have played significant roles as genetic contributors to the modern cultivars. Genome assemblies produced for this study differ in quality, but the estimation of genome assembly and gene annotation quality conducted with BUSCO suggests that they were sufficient to perform comparative analyses. Moreover, we observed that the number of genes grouped in OGs were relatively similar among subspecies, indicating that the potential over-prediction of genes in ‘Maia Oa’ and ‘Calcutta 4’ was mitigated during the clustering procedure. Indeed, over-prediction in draft genomes is expected due to fragmentation, leading to an artefactual increase in the number of genes (Denton et al. 2014).

Although our study is based on one representative per subspecies, *Musa* appears to have a widely shared pangenome, with only a small number of subspecies-specific families identified. The pangenome analysis also reveals a large number of families shared only among subsets of species or subspecies (Figure 1); this “dispensable” genome is thought to contribute to diversity and adaptation (Tettelin et al. 2005; Kahlke et al. 2012). The small number of species-specific OGs in *Musa acuminata* also supports the recent divergence between all genotypes including the split between *M. acuminata* and *M. balbisiana*.

### *M. acuminata* subspecies show a high level of discordance between individual gene trees

By computing gene trees with all single-copy genes OG, we found widespread discordance in gene tree topologies. Topological incongruence can be the result of incomplete lineage sorting, the misassignment of paralogs as orthologs, introgression, or horizontal gene transfer (Maddison 1997). With the continued generation of phylogenomic datasets over the past dozen years, massive amounts of discordance have been reported, first in *Drosophila* (Pollard et al. 2006) and more recently in birds (Jarvis et al. 2014), mammals (Li et al. 2016; Shi & Yang 2018) and plants (Novikova et al. 2016; Pease et al. 2016; Choi et al. 2017; Copetti et al. 2017; Wu et al. 2017). Due to the risk of hemiplasy in such datasets (Avise et al. 2008; Hahn & Nakhleh 2016), we determined that we could not accurately reconstruct either nucleotide substitutions or gene gains and losses among the genomes analyzed here.

In our case, the fact that all possible subspecies tree topologies occurred, and that ratios of minor trees at most nodes were equivalent to those expected under ILS, strongly suggests the presence of ILS (Hahn & Nakhleh 2016). Banana is a paleopolyploid plant that experienced three independent whole genome duplications (WGD), and some fractionation is likely still occurring (D’Hont et al. 2012) (**S. Table 6**). But divergence levels among the single-copy OGs were fairly consistent (Figure 2A), supporting the correct assignment of orthology among sequences. However, we did find evidence for introgression between *malaccensis* and *burmannica*, which contributed a small excess of sites supporting one particular discordant topology (Table 3). This event is also supported by the geographical overlap in the distribution of these two subspecies (Perrier et al. 2011).

The species tree topology supported by all methods used here is different from the tree previously proposed in the literature (**S. Figure 1**). ‘Calcutta 4’ as representative of *M. acuminata* ssp. *burmannica* was placed sister to the other *Musa acuminata* genotypes in our study, whereas several studies have reported proximity between *burmannica* and *malaccensis*, here represented by ‘DH Pahang’ (Janssens et al. 2016; Christelová et al. 2017). Multiple previous studies have attempted to resolve the topology in the *Musa*ceae, but did not include all subspecies considered here, and had very limited numbers of loci. In Christelova et al. (2011), a robust combined approach using maximum likelihood, maximum parsimony, and Bayesian inference was applied to 19 loci, but only *burmannica* and *zebrina* out of the four subspecies were included. Jarret et al. (1992) reported sister relationships between *malaccensis* and *banksii* on the basis of RFLP markers, but did not include any samples from *burmannica* and *zebrina*. It is worth noting that, on the bases of our resolved topology, introgression from *malaccensis* to *burmannica* was detected, and could explain the relationships described previously (Janssens et al. 2016; Sardos et al. 2016).

More strikingly considering previous phylogenetic hypotheses, *malaccensis* appeared most closely related to *banksii*, which is quite distinct from the other *M. acuminata* spp. (Simmonds & Weatherup 1990) and which used to be postulated as its own species based on its geographical area of distribution and floral diversity (Argent 1976). On the bases of genomic similarity, all our analyses support *M. acuminata* ssp. *banksii* as a subspecies of *M. acuminata*.

### Gene tree discordance supports rapid radiation of *Musa acuminata* subspecies

In their evolutionary history, *Musa* species dispersed from ‘northwest to southeast’ into Southeast Asia (Janssens et al. 2016). Due to sea level fluctuations, Malesia (including the nations of Indonesia, Malaysia, Brunei, Singapore, the Philippines, and Papua New Guinea) is a complex geographic region, formed as the result of multiple fusions and subsequent isolation of different islands (Thomas et al. 2012; Janssens et al. 2016). Ancestors of the CalliMusa section (of the *Musa* genus) started to radiate from the northern Indo-Burma region towards the rest of Southeast Asia ~30 MYA, while the ancestors of the Musa (formerly EuMusa/Rhodochlamys) section started to colonize the region ~10 MYA (Janssens et al. 2016). The divergence between *M.acuminata* and *M. balbisiana* has been estimated to be ~5 MYA (Lescot et al. 2008). However, no accurate dating has yet been proposed for the divergence of the *Musa acuminata* subspecies. We hypothesize that after the speciation of *M. acuminata* and *M. balbisiana* (circa 5 MYA) rapid diversification occurred within populations of *M. acuminata*. This hypothesis is consistent with the observed gene tree discordance and high levels of ILS. Such a degree of discordance may reflect a near-instantaneous radiation between all subspecies of *M. acuminata*. Alternatively, it could support the proposed hypothesis of divergence back in the northern part of Malesia during the Pliocene (Janssens et al. 2016), followed by introgression taking place among multiple pairs of species as detected between *malaccensis* and *burmannica*. While massive amounts of introgression can certainly mask the history of lineage splitting (Fontaine et al. 2015), we did not find evidence for such mixing.

Interestingly, such a broad range of gene tree topologies due to ILS (and introgression) has also been observed in gibbons (Carbone et al. 2014; Veeramah et al. 2015; Shi & Yang 2018) for which the area of distribution in tropical forests of Southeast Asia is actually overlapping the center of origin of wild bananas. Moreover, according to Carbone et al. (2014), gibbons also experienced a near-instantaneous radiation ~5 million years ago. It is therefore tempting to hypothesize that ancestors of wild bananas and ancestors of gibbons faced similar geographical isolation and had to colonize and adapt to similar ecological niches, leading to the observed patterns of incomplete lineage sorting.

In this study, we highlighted the phylogenetic complexity in a genome-wide dataset for *Musa acuminata* and *Musa balbisiana*, bringing additional insights to explain why the Musaceae phylogeny has remained controversial. Our work should enable researchers to make inferences about trait evolution, and ultimately should help support crop improvement strategies.

## Material and Methods

### Plant material

Banana leaf samples from accessions ‘Banksii’ (*Musa acuminata* ssp. *banksii*, PT-BA-00024), ‘Maia Oa’ (*Musa acuminata* ssp. *zebrina*, PT-BA-00182) and ‘Calcutta 4’ (*Musa acuminata* ssp. *burmannica*, PT-BA-00051) were supplied by the CRB-Plantes Tropicales Antilles CIRAD-INRA field collection based in Guadeloupe. Leaves were used for DNA extraction. Plant identity was verified at the subspecies level using SSR markers at the *Musa* Genotyping Centre (MGC, Czech Republic) as described in (Christelová, Valarik, et al. 2011) and passport data of the plant is accessible in the *Musa* Germplasm Information System (Ruas et al. 2017). In addition, the representativeness of the genotypes of the four subspecies was verified on a set of 22 samples belonging to the same four *M. acuminata* subspecies of the study (**S. Figure 3**).

### Sequencing and assembly

Genomic DNA was extracted using a modified MATAB method (Risterucci et al. 2000). DNA libraries were constructed and sequenced using the HiSeq2000 (Illumina) technology at BGI (**S. Table 1**). ‘Banksii’ was assembled using SoapDenovo (Luo et al. 2012), and PBJelly2 (English et al. 2012) was used for gap closing using PacBio data generated at the Norwegian Sequencing Center (NSC) with Pacific Biosciences RS II. ‘Maia Oa’ and ‘Calcutta 4’ were assembled using the MaSuRCA assembler (Zimin et al. 2013) (**S. Table 2**). Estimation of genome assembly completeness was assessed with BUSCO plant (Simão et al. 2015) (**S. Table 3**).

### Gene annotation

Gene annotation was performed on the obtained *de novo* assembly for ‘Banksii’, ‘Maia Oa’ and ‘Calcutta 4,’ as well as on the draft *Musa balbisiana* ‘PKW’ assembly (Davey et al, 2013) for consistency and because the published annotation was assessed as low quality. For structural annotation we used EuGene v4.2 (http://eugene.toulouse.inra.fr/) (Foissac et al. 2008) calibrated on *M. acuminata malaccensis* ‘DH Pahang’ reference genome v2, which produced similar results (e.g. number of genes, no missed loci, good specificity and sensitivity) as the official annotation (Martin et al. 2016). EuGene combined genotype-specific (or closely related) transcriptome assemblies, performed with Trinity v2.4 with RNAseq datasets (Sarah et al. 2016), to maximize the likelihood to have genotype-specific gene annotation (**S. Table 4**). The estimation of gene space completeness was assessed with Busco (**S. Table 3**). Because of its high quality and to avoid confusing the community, we did not perform a new annotation for the *M. a. malaccensis* ‘DH Pahang’ reference genome but used the released version 2. Finally, the functional annotation of plant genomes was performed by assigning their associated generic GO terms through the Blast2GO program (Conesa et al. 2005) combining BLAST results from UniProt (E-value 1e-5) (Magrane & Consortium 2011) (**S. Data 1**).

### Gene families

Gene families were identified using OrthoFinder v1.1.4 (Emms & Kelly 2015) with default parameters based on BLASTp (e-value 1e-5). Venn diagrams were made using JVenn online (http://jvenn.toulouse.inra.fr) (Bardou et al. 2014) and alternate visualization was produced with UpsetR (https://gehlenborglab.shinyapps.io/upsetr) (Lex et al. 2014).

### Tree topology from literature

A species tree was initially identified based on previous studies (Sardos et al. 2016; Janssens et al. 2016). Those two studies included all *M. acuminata* subspecies, and had the same tree topology (**S. Figure 1**). In the first study, Sardos et al, (2016) computed a Neighbor-Joining tree from a dissimilarity matrix using bi-allelic GBS-derived SNP markers along the 11 chromosomes of the *Musa* reference genome. Several representatives of each subspecies that comprised genebank accessions related to the genotypes used here were included (Sardos et al. 2016). We annotated the tree to highlight the branches relevant to *M. acuminata* subspecies (**S. Figure 2**). In the second study, a maximum clade credibility tree of Musaceae was proposed based on four gene markers (*rps16, atpB-rbcL, trnL-F* and internal transcribed spacer, ITS) analyzed with Bayesian methods (Janssens et al. 2016).

### Genome-scale phylogenetic analyses and species tree

Single-copy OGs (i.e. orthogroups with one copy of a gene in each of the five genotypes) from protein, coding DNA sequence (CDS), and genes (including introns and UTRs) were aligned with MAFFT v7.271 (Katoh & Standley 2013), and gene trees were constructed using PhyML v3.1 (Guindon et al. 2009) with ALrT branch support. All trees were rooted using *Musa balbisiana* as outgroup using Newick utilities v1.6 (Junier & Zdobnov 2010) (**S. Data 2**). Individual gene tree topologies were visualized as a cloudogram with DensiTree v2.2.5 (Bouckaert 2010).

Single-copy OGs were further investigated with the quartet method implemented in ASTRAL v5.5.6 (Mirarab & Warnow 2015). In parallel, we carried out a Supertree approach following the SSIMUL procedure (http://www.atgc-montpellier.fr/ssimul/) (Scornavacca et al. 2011) combined with PhySIC_IST (http://www.atgc-montpellier.fr/physic_ist/) (Scornavacca et al. 2008) applied to a set of rooted trees corresponding to core OGs (including single and multiple copies), and accessory genes for which only one representative species was missing (except outgroup species). Finally, single-copy OGs (CDS only) were used to generate a concatenated genome-scale alignment with FASconCAT-G (Kück & Longo 2014) and a tree was constructed using PhyML (NNI, HKY85, 100 bootstrap).

### Search for introgression

Ancient gene flow was assessed with the ABBA-BABA test or *D*-statistic (Green et al. 2010; Durand et al. 2011) and computed on the concatenated multiple alignment converted to the MVF format and processed with MVFtools (Pease & Rosenzweig 2017), similar to what is described in Wu et al. (2017) (https://github.com/wum5/JaltPhylo). The direction of introgression was further assessed with the *D*_2_ test (Hibbins and Hahn, unpublished). The *D*_2_ statistic captures differences in the heights of genealogies produced by introgression occurring in alternate directions by measuring the average divergence between species A and C in gene trees with an ((A,B),C) topology (denoted [d_AC_|A,B]), and subtracting the average A-C divergence in gene trees with a ((B,C),A) topology (denoted [d_AC_|B,C]), so that *D*_2_ = (d_AC_|A,B) - (d_AC_|B,C). If the statistic is significantly positive, it means that introgression has either occurred in the B→C direction or in both directions. *D*_2_ significance was assessed by permuting labels on gene trees 1000 times and calculating *P*-values from the resulting null distribution of *D*_2_ values. The test was implemented with a Perl script using distmat from EMBOSS (Rice et al. 2000) with Tajima-Nei distance applied to multiple alignments associated with gene trees fitting the defined topologies above (https://github.com/mrouard/perl-script-utils).

## Data availability

Raw sequence reads for *de novo* assemblies were deposited in the Sequence Read Archive (SRA) of the National Center for Biotechnology Information (NCBI) (BioProject: PRJNA437930 and SRA: SRP140622). Assembly and gene annotation data are available on the Banana Genome Hub (Droc et al, 2013) (http://banana-genome-hub.southgreen.fr/species-list). Cluster and gene tree results are available on a dedicated database (http://panMusa.greenph.org) hosted on the South Green Bioinformatics Platform (Guignon et al. 2016).

## Acknowledgments

We thank Noel Chen and Qiongzhi He (BGI) for providing sequencing services with Illumina and Ave Tooming-Klunderud (CEES) for PacBio sequencing services and Computomics for support with assembly. We thank Erika Sallet (INRA) for providing early access to the new version of Eugene with helpful suggestions. We thank the CRB-Plantes Tropicales Antilles CIRAD-INRA for providing plant materials. We would like also to acknowledge Jae Young Choi (NYU), Steven Janssens (MBG), Laura Kubatko (OSU) for helpful discussions and advice on species tree topologies. This work was financially supported by CGIAR Fund Donors and CGIAR Research Programme on Roots, Tubers and Bananas (RTB) and technically supported by the high performance cluster of the UMR AGAP – CIRAD of the South Green Bioinformatics Platform (http://www.southgreen.fr).

## Authors contribution

MR, NR and AD set up the study and MR coordinated the study. AD and FCB provided access to plant material and DNA. NY provided access to transcriptome data and GM to repeats library for gene annotation. BG performed assembly and gap closing. MR, GD, GM, YH, JS and AC performed analyses. VB, MSH, and MWH provided guidance on methods and helped with result interpretation. VG and MR set up the PanMusa website. MR wrote the manuscript with significant contributions from MWH, VB, and JS, and all co-authors commented on the manuscript

## Additional information

**S. Figure 1. Species tree of Musa acuminata subspecies extrapolated from literature review**

**S. Figure 2. Neighbor-Joining tree from 105 M. acuminata and cultivated accessions**

**S. Figure 3. Individual ancestries investigated with the Admixture software package**

**S. Table 1. Libraries used for the genome assemblies**

**S Table 2. Summary of the genome assembly**

**S. Table 3. Results of gene space assessment with BUSCO**

**S. Table 4. Summary of the genome annotation**

**S. Table 5. Global summary of the gene clustering**

**S. Table 6. List of 18 phylogenetic informative shared single copy nuclear genes from Duarte et al. 2010 mapped to *Musa* genomes.**

**S. Data 1. List of Gene ontology mapped by genomes**

**S. Data 2. List of gene trees obtained at protein-coding, CDS and gene based level**

## References

Alexander DH, Lange K. 2011. Enhancements to the ADMIXTURE algorithm for individual ancestry estimation. BMC Bioinformatics. 12:246. doi: 10.1186/1471-2105-12-246.

Argent G. 1976. The wild bananas of Papua New Guinea. Notes Roy Bot Gard Edinb. 35:77–114.

Avise JC, Robinson TJ, Kubatko L. 2008. Hemiplasy: A New Term in the Lexicon of Phylogenetics. Syst. Biol. 57:503–507. doi: 10.1080/10635150802164587.

Bardou P, Mariette J, Escudié F, Djemiel C, Klopp C. 2014. jvenn: an interactive Venn diagram viewer. BMC Bioinformatics. 15:293. doi: 10.1186/1471-2105-15-293.

Bouckaert RR. 2010. DensiTree: making sense of sets of phylogenetic trees. Bioinformatics. 26:1372–1373. doi: 10.1093/bioinformatics/btq110.

Bravo GA et al. 2018. Embracing heterogeneity: Building the Tree of Life and the future of phylogenomics. PeerJ Inc. doi: 10.7287/peerj.preprints.26449v3.

Carbone L et al. 2014. Gibbon genome and the fast karyotype evolution of small apes. Nature. 513:195–201. doi: 10.1038/nature13679.

Cheesman EE. 1948. Classification of the Bananas. Kew Bull. 3:17–28. doi: 10.2307/4118909.

Choi JY et al. 2017. The Rice Paradox: Multiple Origins but Single Domestication in Asian Rice. Mol. Biol. Evol. 34:969–979. doi: 10.1093/molbev/msx049.

Christelová P, Valarik M, et al. 2011. A platform for efficient genotyping in Musa using microsatellite markers. AoB Plants. 2011:plr024–plr024. doi: 10.1093/aobpla/plr024.

Christelová P et al. 2017. Molecular and cytological characterization of the global Musa germplasm collection provides insights into the treasure of banana diversity. Biodivers. Conserv. 26:801–824. doi: 10.1007/s10531-016-1273-9.

Christelová P, Valárik M, Hřibová E, De Langhe E, Doležel J. 2011. A multi gene sequence-based phylogeny of the Musaceae (banana) family. BMC Evol. Biol. 11:103. doi: 10.1186/14712148-11-103.

Conesa A et al. 2005. Blast2GO: a universal tool for annotation, visualization and analysis in functional genomics research. Bioinforma. Oxf. Engl. 21:3674–3676. doi: 10.1093/bioinformatics/bti 610.

Copetti D et al. 2017. Extensive gene tree discordance and hemiplasy shaped the genomes of North American columnar cacti. Proc. Natl. Acad. Sci. 114:12003–12008. doi: 10.1073/pnas.1706367114.

Dagan T, Martin W. 2006. The tree of one percent. Genome Biol. 7:118. doi: 10.1186/gb-20067-10-118.

Davey MW et al. 2013. A draft Musa balbisiana genome sequence for molecular genetics in polyploid, inter- and intra-specific Musa hybrids. BMC Genomics. 14:683. doi: 10.1186/14712164-14-683.

De Langhe E et al. 2009. Why Bananas Matter: An introduction to the history of banana domestication. Ethnobot. Res. Appl. 7:165–177. doi: 10.17348/era.7.0.165-177.

Denton JF et al. 2014. Extensive Error in the Number of Genes Inferred from Draft Genome Assemblies. PLOS Comput. Biol. 10:e1003998. doi: 10.1371/journal.pcbi.1003998.

D’Hont A et al. 2012. The banana (Musa acuminata) genome and the evolution of monocotyledonous plants. Nature. doi: 10.1038/nature11241.

Durand EY, Patterson N, Reich D, Slatkin M. 2011. Testing for Ancient Admixture between Closely Related Populations. Mol. Biol. Evol. 28:2239–2252. doi: 10.1093/molbev/msr048.

Emms DM, Kelly S. 2015. OrthoFinder: solving fundamental biases in whole genome comparisons dramatically improves orthogroup inference accuracy. Genome Biol. 16:157. doi: 10.1186/s13059-015-0721-2.

English AC et al. 2012. Mind the Gap: Upgrading Genomes with Pacific Biosciences RS Long-Read Sequencing Technology. PLOS ONE. 7:e47768. doi: 10.1371/journal.pone.0047768.

Foissac S et al. 2008. Genome Annotation in Plants and Fungi: EuGene as a Model Platform. Curr. Bioinforma. http://www.eurekaselect.com/82677/article (Accessed March 1, 2018).

Folk Ryan A., Soltis Pamela S., Soltis Douglas E., Guralnick Robert. 2018. New prospects in the detection and comparative analysis of hybridization in the tree of life. Am. J. Bot. 0. doi: 10.1002/ajb2.1018.

Fontaine MC et al. 2015. Extensive introgression in a malaria vector species complex revealed by phylogenomics. Science. 347:1258524. doi: 10.1126/science.1258524.

Guignon V et al. 2016. The South Green portal: A comprehensive resource for tropical and Mediterranean crop genomics. Curr. Plant Biol. 7:6–9.

Guindon S, Delsuc F, Dufayard J-F, Gascuel O. 2009. Estimating maximum likelihood phylogenies with PhyML. Methods Mol. Biol. Clifton NJ. 537:113–137. doi: 10.1007/978-1-59745-251-9_6.

Hahn MW, Nakhleh L. 2016. Irrational exuberance for resolved species trees. Evol. Int. J. Org. Evol. 70:7–17. doi: 10.1111/evo.12832.

Janssens SB et al. 2016. Evolutionary dynamics and biogeography of Musaceae reveal a correlation between the diversification of the banana family and the geological and climatic history of Southeast Asia. New Phytol. 210:1453–1465. doi: 10.1111/nph.13856.

Jarret R, Gawel N, Whittemore A, Sharrock S. 1992. RFLP-based phylogeny of Musa species in Papua New Guinea. Theor. Appl. Genet. 84–84. doi: 10.1007/BF00224155.

Jarvis ED et al. 2014. Whole-genome analyses resolve early branches in the tree of life of modern birds. Science. 346:1320–1331. doi: 10.1126/science.1253451.

Junier T, Zdobnov EM. 2010. The Newick utilities: high-throughput phylogenetic tree processing in the UNIX shell. Bioinforma. Oxf. Engl. 26:1669–1670. doi: 10.1093/bioinformatics/btq243.

Katoh K, Standley DM. 2013. MAFFT multiple sequence alignment software version 7: improvements in performance and usability. Mol. Biol. Evol. 30:772–780. doi: 10.1093/molbev/mst010.

Kreft L, Botzki A, Coppens F, Vandepoele K, Van Bel M. 2017. PhyD3: a phylogenetic tree viewer with extended phyloXML support for functional genomics data visualization. Bioinforma. Oxf. Engl. doi: 10.1093/bioinformatics/btx324.

Kück P, Longo GC. 2014. FASconCAT-G: extensive functions for multiple sequence alignment preparations concerning phylogenetic studies. Front. Zool. 11:81. doi: 10.1186/s12983-014-0081-x.

Lescot M et al. 2008. Insights into the Musa genome: Syntenic relationships to rice and between Musa species. BMC Genomics. 9:58. doi: 10.1186/1471-2164-9-58.

Lex A, Gehlenborg N, Strobelt H, Vuillemot R, Pfister H. 2014. UpSet: Visualization of Intersecting Sets. IEEE Trans. Vis. Comput. Graph. 20:1983–1992. doi: 10.1109/TVCG.2014.2346248.

Li G, Davis BW, Eizirik E, Murphy WJ. 2016. Phylogenomic evidence for ancient hybridization in the genomes of living cats (Felidae). Genome Res. 26:1–11. doi: 10.1101/gr.186668.114.

Luo R et al. 2012. SOAPdenovo2: an empirically improved memory-efficient short-read de novo assembler. GigaScience. 1:18. doi: 10.1186/2047-217X-1-18.

Maddison WP. 1997. Gene Trees in Species Trees. Syst. Biol. 46:523–536. doi: 10.1093/sysbio/46.3.523.

Magrane M, Consortium U. 2011. UniProt Knowledgebase: a hub of integrated protein data. Database J. Biol. Databases Curation. 2011. doi: 10.1093/database/bar009.

Martin G et al. 2016. Improvement of the banana “Musa acuminata” reference sequence using NGS data and semi-automated bioinformatics methods. BMC Genomics. 17:243. doi: 10.1186/s12864-016-2579-4.

Medini D, Donati C, Tettelin H, Masignani V, Rappuoli R. 2005. The microbial pan-genome. Curr. Opin. Genet. Dev. 15:589–594. doi: 10.1016/j.gde.2005.09.006.

Mirarab S, Warnow T. 2015. ASTRAL-II: coalescent-based species tree estimation with many hundreds of taxa and thousands of genes. Bioinforma. Oxf. Engl. 31:i44–52. doi: 10.1093/bioinformatics/btv234.

Morgante M, De Paoli E, Radovic S. 2007. Transposable elements and the plant pan-genomes. Curr. Opin. Plant Biol. 10:149–155. doi: 10.1016/j.pbi.2007.02.001.

Novikova PY et al. 2016. Sequencing of the genus Arabidopsis identifies a complex history of nonbifurcating speciation and abundant trans-specific polymorphism. Nat. Genet. 48:1077–1082. doi: 10.1038/ng.3617.

Pease J, Rosenzweig B. 2017. Encoding Data Using Biological Principles: the Multisample Variant Format for Phylogenomics and Population Genomics. IEEE/ACM Trans. Comput. Biol. Bioinform. PP:1–1. doi: 10.1109/TCBB.2015.2509997.

Pease JB, Haak DC, Hahn MW, Moyle LC. 2016. Phylogenomics Reveals Three Sources of Adaptive Variation during a Rapid Radiation. PLOS Biol. 14:e1002379. doi: 10.1371/journal.pbio.1002379.

Perrier X et al. 2011. Multidisciplinary perspectives on banana (Musa spp.) domestication. Proc. Natl. Acad. Sci. doi: 10.1073/pnas.1102001108.

Pollard DA, Iyer VN, Moses AM, Eisen MB. 2006. Widespread Discordance of Gene Trees with Species Tree in Drosophila: Evidence for Incomplete Lineage Sorting. PLOS Genet. 2:e173. doi: 10.1371/journal.pgen.0020173.

Rice P, Longden I, Bleasby A. 2000. EMBOSS: the European Molecular Biology Open Software Suite. Trends Genet. TIG. 16:276–277.

Risterucci AM et al. 2000. A high-density linkage map of *Theobroma cacao* L. Theor. Appl. Genet. 101:948–955. doi: 10.1007/s001220051566.

Rouard M et al. 2011. GreenPhylDB v2.0: comparative and functional genomics in plants. Nucleic Acids Res. 39:D1095–1102. doi: 10.1093/nar/gkq811.

Ruas M et al. 2017. MGIS: managing banana (Musa spp.) genetic resources information and high-throughput genotyping data. Database. 2017. doi: 10.1093/database/bax046.

Sarah G et al. 2016. A large set of 26 new reference transcriptomes dedicated to comparative population genomics in crops and wild relatives. Mol. Ecol. Resour. doi: 10.1111/17550998.12587.

Sardos Julie et al. 2016. A Genome-Wide Association Study on the Seedless Phenotype in Banana (Musa spp.) Reveals the Potential of a Selected Panel to Detect Candidate Genes in a Vegetatively Propagated Crop. PLOS ONE. 11:e0154448. doi: 10.1371/journal.pone.0154448.

Sardos J. et al. 2016. DArT whole genome profiling provides insights on the evolution and taxonomy of edible Banana (Musa spp.). Ann. Bot. mcw170. doi: 10.1093/aob/mcw170.

Scornavacca C, Berry V, Lefort V, Douzery EJ, Ranwez V. 2008. PhySIC_IST: cleaning source trees to infer more informative supertrees. BMC Bioinformatics. 9:413. doi: 10.1186/1471-21059-413.

Scornavacca C, Berry V, Ranwez V. 2011. Building species trees from larger parts of phylogenomic databases. Inf. Comput. 209:590–605. doi: 10.1016/j.ic.2010.11.022.

Shi C-M, Yang Z. 2017. Coalescent-based analyses of genomic sequence data provide a robust resolution of phylogenetic relationships among major groups of gibbons. Mol. Biol. Evol. doi: 10.1093/molbev/msx277.

Shi C-M, Yang Z. 2018. Coalescent-Based Analyses of Genomic Sequence Data Provide a Robust Resolution of Phylogenetic Relationships among Major Groups of Gibbons. Mol. Biol. Evol. 35:159–179. doi: 10.1093/molbev/msx277.

Simão FA, Waterhouse RM, Ioannidis P, Kriventseva EV, Zdobnov EM. 2015. BUSCO: assessing genome assembly and annotation completeness with single-copy orthologs. Bioinformatics. 31:3210–3212. doi: 10.1093/bioinformatics/btv351.

Simmonds NW. 1956. Botanical Results of the Banana Collecting Expedition, 1954-5. Kew Bull. 11:463–489. doi: 10.2307/4109131.

Simmonds NW. 1962. The evolution of the bananas. Longmans: London (GBR).

Simmonds NW, Shepherd K. 1955. The taxonomy and origins of the cultivated bananas. J. Linn. Soc. Lond. Bot. 55:302–312. doi: 10.1111/j.1095-8339.1955.tb00015.x.

Simmonds NW, Weatherup STC. 1990. Numerical taxonomy of the wild bananas (Musa). New Phytol. 115:567–571. doi: 10.1111/j.1469-8137.1990.tb00485.x.

Tettelin H et al. 2005. Genome analysis of multiple pathogenic isolates of Streptococcus agalactiae: Implications for the microbial “pan-genome”. Proc. Natl. Acad. Sci. U. S. A. 102:13950–13955. doi: 10.1073/pnas.0506758102.

Thomas DC et al. 2012. West to east dispersal and subsequent rapid diversification of the mega-diverse genus Begonia (Begoniaceae) in the Malesian archipelago. J. Biogeogr. 39:98–113. doi: 10.1111/j. 1365-2699.2011.02596.x.

Veeramah KR et al. 2015. Examining Phylogenetic Relationships Among Gibbon Genera Using Whole Genome Sequence Data Using an Approximate Bayesian Computation Approach. Genetics. 200:295–308. doi: 10.1534/genetics.115.174425.

Wu M, Kostyun JL, Hahn MW, Moyle L. 2017. Dissecting the basis of novel trait evolution in a radiation with widespread phylogenetic discordance. bioRxiv. 201376. doi: 10.1101/201376.

Yachdav G et al. 2016. MSAViewer: interactive JavaScript visualization of multiple sequence alignments. Bioinforma. Oxf. Engl. 32:3501–3503. doi: 10.1093/bioinformatics/btw474.

Zimin AV et al. 2013. The MaSuRCA genome assembler. Bioinformatics. 29:2669–2677. doi: 10.1093/bioinformatics/btt476.

